# Sexual Dimorphic Metabolic and Cognitive Responses of C57BL/6 Mice to Fisetin or Dasatinib and Quercetin Cocktail Oral Treatment

**DOI:** 10.1101/2021.11.08.467509

**Authors:** Yimin Fang, David Medina, Robert Stockwell, Sam McFadden, Kathleen Quinn, Mackenzie R. Peck, Andrzej Bartke, Kevin N. Hascup, Erin R. Hascup

## Abstract

Senolytic treatment in aged mice clears senescent cell burden leading to functional improvements. However, less is known regarding the effects of these compounds when administered prior to significant senescent cell accumulation. From 4-13 months of age, C57BL/6 male and female mice received monthly oral dosing of either 100 mg/kg Fisetin or a 5 mg/kg Dasatinib (D) plus 50 mg/kg Quercetin (Q) cocktail. During treatment, several aspects of healthy aging were assayed including glucose metabolism using an insulin and glucose tolerance test, cognitive performance using Morris water maze and novel object recognition, and energy metabolism using indirect calorimetry. Afterwards, mice were euthanized for plasma and tissue specific markers of senescence-associated secretory phenotype (SASP) and white adipose tissue accumulation (WAT). Sexually dimorphic treatment effects were observed. Fisetin treated male mice had reduced SASP, enhanced glucose and energy metabolism, improved cognitive performance, and increased mRNA expression of adiponectin receptor 1 and glucose transporter 4. D+Q treatment had minimal effects in male C57BL/6 mice, but was detrimental to females causing increased SASP expression along with accumulation of WAT depots. Reduced energy metabolism and cognitive performance were also noted. Fisetin treatment had no effect in female C57BL/6 mice potentially due to a slower rate of biological aging. In summary, the senolytic treatment in young adulthood, has beneficial, negligible, or detrimental effects in C57BL/6 mice dependent upon sex and treatment. These observations should serve as a note of caution in this rapidly evolving and expanding field of investigation.

**Graphical Abstract:** Male and female C57BL/6 mice were treated with once monthly oral doses of either Dasatinib (D) + Quercetin (Q) or Fisetin from 4-13 months of age. Females treated with D+Q had increased adiposity and SASP markers (red spheres) along with decreased metabolism (blue flame) and cognitive performance. Males treated with Fisetin had reduced SASP markers (blue spheres) as well as improved metabolism (red flame) and cognition. No effects were observed in females treated with Fisetin or males treated with D+Q.

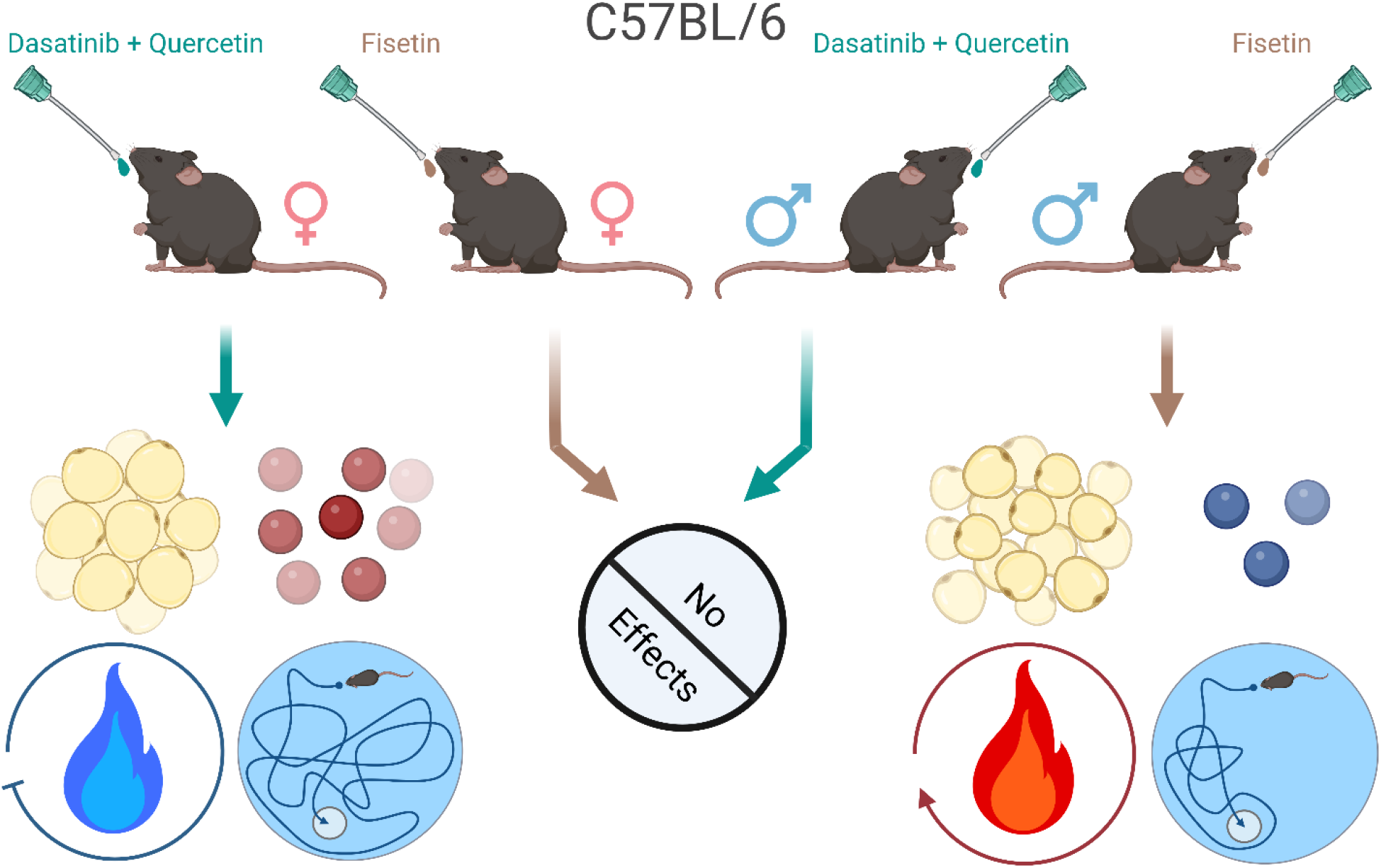

## Introduction

Fisetin and quercetin (Q) are plant-derived flavonoids that offer cytoprotection against cellular stress and act as anti-inflammatory, chemopreventive, chemotherapeutic, and senotherapeutic agents (1, 2). Additionally, Q incites immune response against allergic diseases (2, 3). Dasatinib (D) is a tyrosine kinase inhibitor used to treat leukemia (4) and is routinely used in combination with Q to improve the senotherapeutic potency. Although both prooxidan flavonoids are structurally analogous, Fisetin has higher senolytic activity than Q whose potency is also reliant on endogenous trace elements such as iron and copper (5).

As the first senolytic drugs, they were discovered to selectively clear senescent cells generated during cellular senescence (6). Cell senescence is a phenomenon defined as a stable cell cycle arrest, and is associated with fundamental aging processes and age-related diseases (7). Although this process is essential for preventing replication of damaged DNA thereby suppressing tumor formation for cancer, accumulation of senescent cells with aging produces senescence-associated secretory phenotype (SASP) (7, 8). SASP includes proinflammatory cytokines, chemokines, growth factors, and proteases (9, 10) caused by chronic inflammation (10-12), DNA damage (13, 14), mitochondrial dysfunction (15), immune cell dysfunction (16), ROS generation (13, 17), and brain protein aggregation (18, 19). All of which may be factors that predispose individuals to a multitude of age-related disorders (7).

Fisetin and D+Q selectively clear senescent cells (6), thereby delaying aging-associated disorders and improving healthspan and lifespan. This has been observed after reducing senescent cell burden in progeroid or aged (twenty-four-month-old) C57BL/6 mice (20-22). Moreover, deletion of senescent cells from the brain genetically or pharmacologically with senolytic drugs led to functional improvements in mouse models of neurodegenerative diseases such as Parkinson’s and Alzheimer’s disease (18, 23-25). These studies have shown senotherapeutics can reduce senescent cell burden and have positive impacts on animals with accelerated aging, advanced age, or neurodegenerative disorders. Accordingly, senotherapeutics are currently marketed as anti-aging therapies where young, healthy adults can take these products as dietary supplements.

However, less is known about the effects of these compounds when administered prior to significant senescent cell accumulation. Thus, the experiments were designed to examine the long-term effects of monthly oral Fisetin or D+Q treatment when initiated in young (four-month-old) C57BL/6 mice. We examined morphological, metabolic, physical, and cognitive components that are known to be affected by senescent cell accumulation. The results presented here indicate that monthly administration of Fisetin or D+Q had sexually dimorphic effects which also depended on treatment type in C57BL/6 mice. We highlight a potential new mechanism of action involving the beneficial roles of glucose and adiponectin signaling both peripherally and centrally by early and long-term administration of Fisetin on male C57BL/6 mice.

## Results

### Administration of Fisetin or D+Q altered the SASP profile of C57BL/6 mice in a sex dependent manner

It has been reported that senolytic drug treatment decreased senescent burden in twenty-four-month-old C57BL/6 mice leading to functional improvements (20). However, it was unknown whether the same treatment administered monthly initiated during young adulthood (four months of age) would improve the overall health of C57BL/6 mice as they aged. To examine this, Fisetin or a cocktail of D+Q were orally administered in C57BL/6 mice starting at four months of age and continued once each month for nine months. All mice underwent glucose tolerance test (GTT) and insulin tolerance test (ITT) to evaluate glucose homeostasis, indirect calorimetry to determine energy metabolism, grip force to monitor muscle strength, and Morris water maze (MWM) and novel object recognition (NOR) to ascertain cognitive abilities. At approximately thirteen months of age, the mice were sacrificed to determine body composition and collect plasma and tissues for analyses of SASP markers, lipid metabolism, glucose and adiponectin signaling, and synaptic plasticity. The experimental paradigm is shown in Fig 1a.

**Figure 1:**
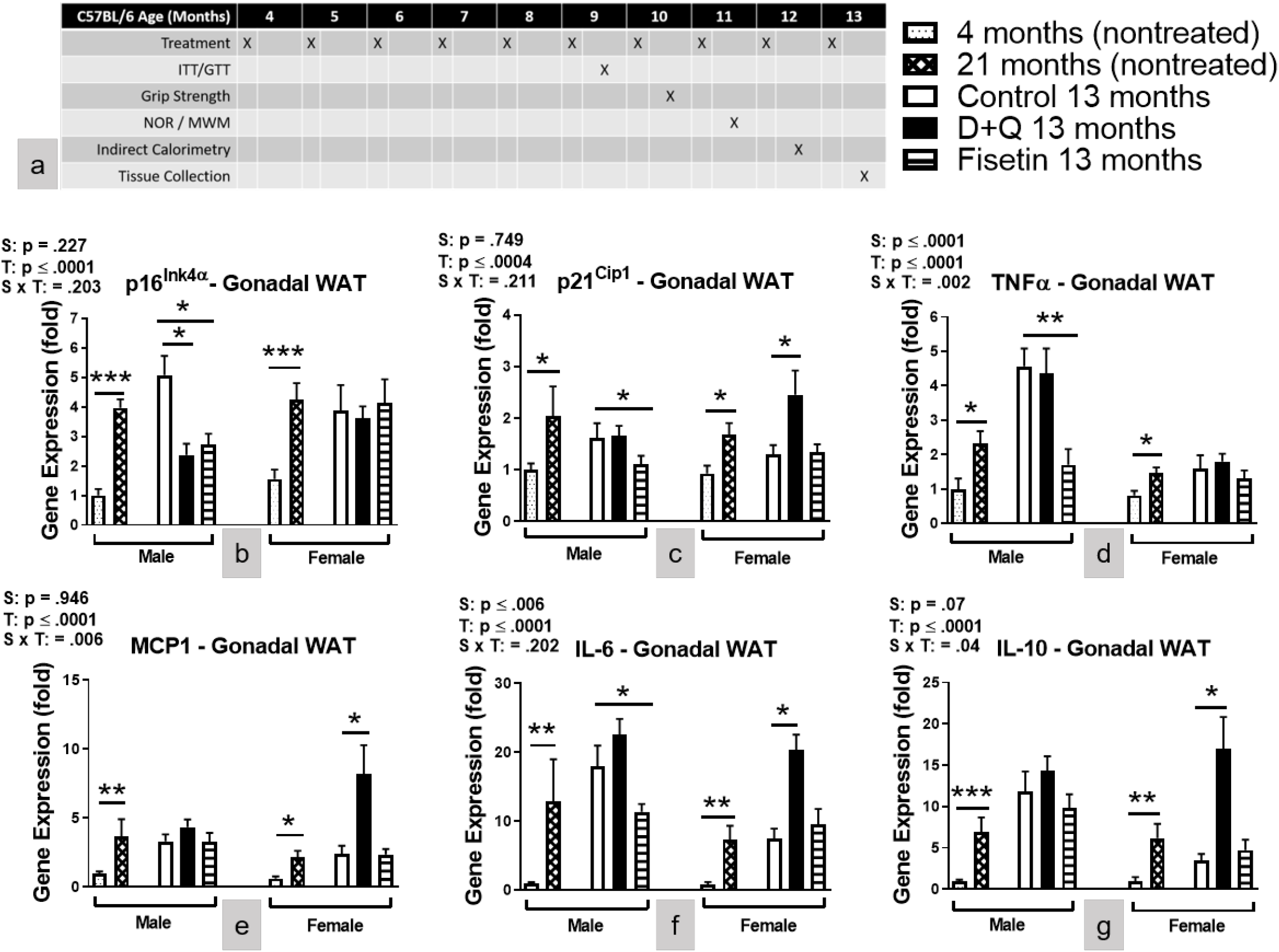
Senotherapeutic treatment altered the SASP profile in C57BL/6 mice. (a) Experimental paradigm. (b-g) Gene expressions in gonadal fat depot. Data are represented as means ± *SEM* (*n* = 8–12). A two-way ANOVA was used to determine P-values for the categorial variables (S = Sex and T = Treatment) and their interaction (S x T), which are shown for each graph. \**p* ≤ 0.05, \*\**p* ≤ 0.01, ***p ≤ 0.001 based on a two-tailed Student’s *t* test.

To investigate whether the compounds used in this study decreased SASP profiles and other senescent markers, the gene expressions were measured in gonadal white adipose tissue (WAT) by RT-PCR. To evaluate aging effects, additional groups of treatment naïve young (four months of age) and old (twenty-one months of age) mice were studied. We assayed for SASP markers including p16^Ink4α^, p21^Cip1^, TNFα, MCP 1, IL-6, and IL-10. Although IL-10 is a potent anti-inflammatory cytokine, studies have shown IL-10 activates cellular senescence mechanisms (26) and is secreted from senescent cells (27). The results showed that in both the male and female mice, the transcriptional levels of these SASP markers were lower in the treatment naïve young compared to old C57BL/6 mice (Fig. 1b-g). The vehicle treated littermate control C57BL/6 mice sacrificed at thirteen months of age had similar mRNA levels of the above mentioned SASP markers as the untreated twenty-one-month-old mice (Fig. 1b-g). This is consistent with recent tissue and age-specific RNA sequencing in both sexes of C57BL/6 mice where a significant increase in differentially expressed genes was first evident in gonadal WAT that occurred by mid-age and was maintained at ages older than twenty months (28). Others have shown p16^Ink4α^ and p21^Cip1^ have similar mRNA expression levels at 12 and 24 months of age in the hypothalamus, heart, liver, and kidney of female CB6F1 mice (29).

We also observed that C57BL/6 males had higher levels of mRNA expression of TNFα and IL-6 (Fig. 1d, f) compared to littermate females. Interestingly, the effects of the treatments were sex-dependent. The treatment with Fisetin or D+Q reduced mRNA levels of p16^Ink4α^ in the male mice, but had no effects in the female mice (Fig. 1b). The treatment with Fisetin reduced the mRNA levels of p21^Cip1^, TNFα, and IL-6 in the male mice, but had no effect in the female mice (Figs. 1c, 1d, & 1f), while the treatment with D+Q raised the mRNA levels of p21^Cip1^, MCP1, IL-6, and IL-10 (Figs. 1c & 1e-g) in the female mice.

The same SASP mRNA levels were examined in the hippocampus (SI Fig. 1a-f). mRNA levels were higher in both sexes of treatment naïve twenty-one-month-old C57BL/6 mice compared to four-month-old littermates. p21^Cip1^ was the only SASP marker increased in the hippocampus of female C57BL/6 mice compared to male littermates. In general, mRNA hippocampal expression patterns after senotherapeutic treatment was similar to observations in gonadal WAT. In male C57BL/6 mice, Fisetin treatment reduced mRNA levels of p16^Ink4α^ and p21^Cip1^ (SI Fig 1a-b), but did not affect the other assayed cytokines (Fig 1c-f). Reduced hippocampal mRNA levels of p21^Cip1^ (SI Fig. 1b) were also observed in female C57BL/6 mice. D+Q treatment increased mRNA levels of TNFα (SI Fig. 1c) and IL-10 (SI Fig. 1e) in both sexes of C57BL/6 mice.

Plasma concentrations of TNFα, MCP1, IL-6, and IL-10 (SI Fig. 2a-d) were elevated in both sexes of treatment naïve twenty-one-month-old C57BL/6 mice compared to four-month-old littermates. The plasma concentration of TNFα was increased in males, while IL-10 was increased in females compared to sex-matched C57BL/6 littermates. Fisetin treatment decreased the concentration of MCP1 (SI Fig. 2b) in males while increasing IL-10 (SI Fig. 2d) concentration in female C57BL/6 mice. D+Q treatment had no effect in males, but increased the concentration of MCP1, IL-6, and IL-10 (SI Fig. b-d) in female C57BL/6 mice.

Altogether, the data showed that Fisetin treatment reduced the levels of SASP markers in the plasma as well as the peripheral and central tissue of male C57BL/6 mice, while little to no changes were observed in female littermates. The D+Q treatment increased, rather than decreased, SASP markers in female C57BL/6 mice. In male C57BL/6 mice, D+Q treatment had minimal to no effects in plasma and WAT, but increased hippocampal TNFα mRNA expression.

### D+Q treatment led to increased body weight and adiposity in female C57BL/6 mice while Fisetin treatment improved glucose metabolism and increased plasma adiponectin in male C57BL/6 mice

Given the observation that C57BL/6 mice responded to the treatment in a sexually dimorphic manner, it was important to know how the physiological parameters including body composition, glucose utilization, and energy metabolism were affected by senotherapeutic treatment. The results showed that body weight in the males was not changed by either treatment, while it was increased in the female C57BL/6 mice receiving monthly D+Q administration (Fig. 2a-b). Body composition analysis indicated that the increased body weight of D+Q treated female C57BL/6 mice was attributed to an elevation of adiposity including subcutaneous (SC) and gonadal WAT depots (Figs. 2c-e). No changes in body weight or composition were observed after nine months of Fisetin treatment in female C57BL/6 mice, and no changes in these parameters were detected in male C57BL/6 mice receiving either of the treatments (Figs 2a-g). Body composition also showed sex differences whereby female C57BL/6 mice had a higher percentage of SC and gonadal WAT while males had a larger perirenal depot (Fig. 2d-f).

**Figure 2:**
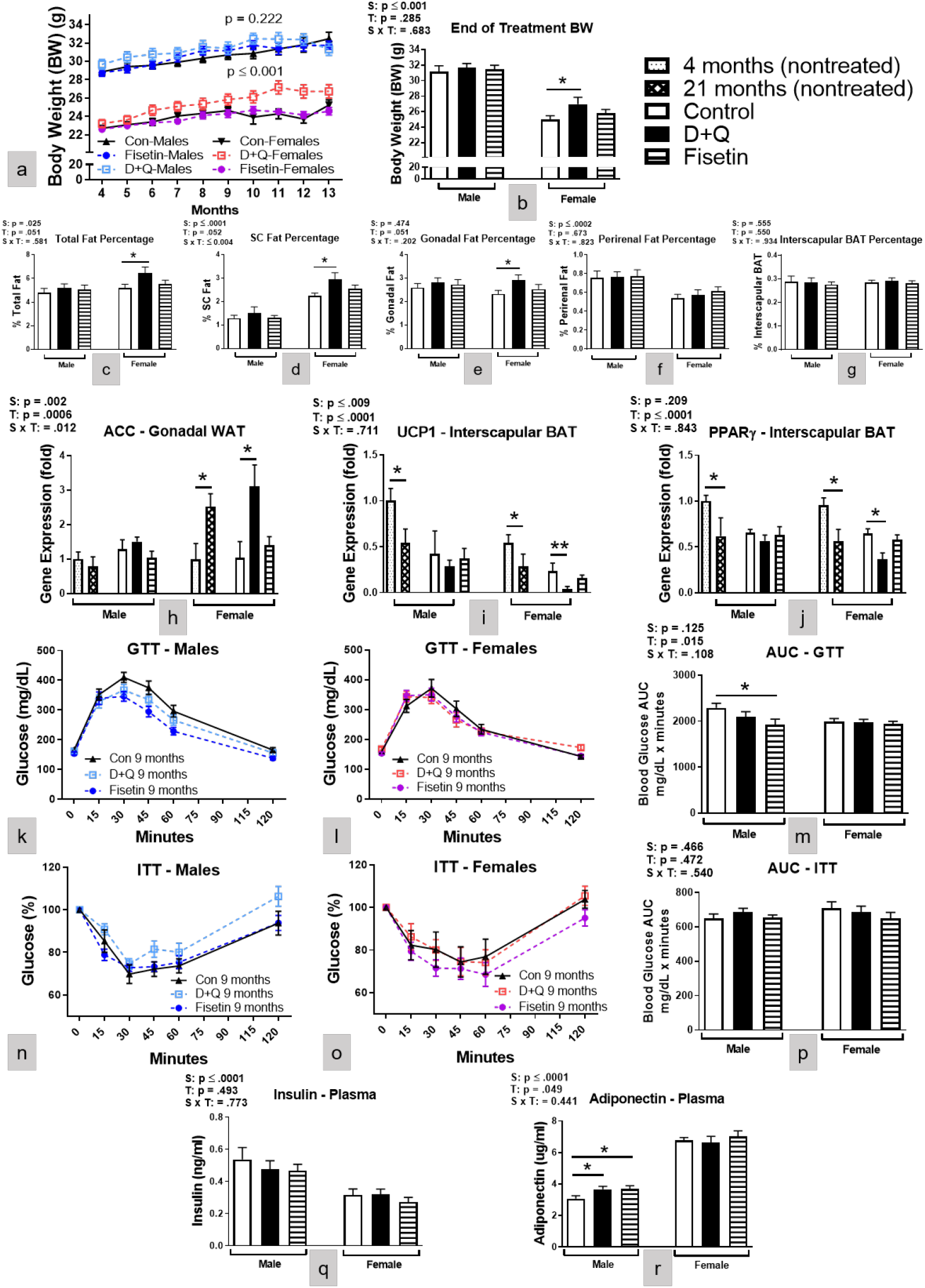
D+Q treatment increased abdominal WAT in female mice while Fisetin treatment improved glucose metabolism and increased plasma adiponectin in male mice. Mouse body weight (a-b, BW) along with percentage of total (c) and different adipose tissue depots present in relation to BW (d-g). Gene expression of ACC in gonadal WAT (h) as well as UCP1 and PPARγ in interscapular BAT (i-j) depots. Glucose tolerance test (GTT), insulin tolerance test (ITT), and their respective area under the curve (AUC) after five treatments (k-p). Plasma insulin and adiponectin concentrations (q-r) taken at time of euthanization after 10 treatments. Data are means ± SEM (n = 8–20). A two-way ANOVA was used to determine P-values for the categorial variables (S = Sex and T = Treatment) and their interaction (S x T), which are shown for each bar graph. *p ≤ 0.05, **p ≤ 0.01 based on a two-tailed Student’s t test.

The accumulation of WAT in D+Q treated female mice could result from upregulated lipid deposition or downregulated thermogenic activity in brown adipose tissue (BAT). Thus, the expressions of genes related to lipid metabolism in gonadal WAT, and to uncoupling of the mitochondrial electron transport chain in BAT were examined. The results showed that Acetyl-CoA carboxylase (ACC), the rate-limiting enzyme for lipid synthesis, was up-regulated in gonadal WAT of D+Q treated female C57BL/6 mice (Fig. 2h). In contrast to WAT, uncoupling protein 1 (UCP1), a key regulator of non-shivering thermogenesis, and its upstream activator PPARγ (30, 31) were downregulated in BAT of D+Q treated female mice (Fig. 2i-j). Sex effects were observed with higher ACC in female C57BL/6 gonadal WAT and UCP1 in male interscapular BAT. The data from WAT and BAT support that D+Q treated female mice had higher lipid synthesis activity, mediated by increased ACC, and lower thermogenic activity, via reduced UCP1 and PPARγ. These changes could cause WAT accumulation subsequently leading to the observed SASP increases after D+Q treatment in female C57BL/6 mice.

Considering adipocytes are involved in glucose metabolism, we wanted to determine treatment effects on peripheral glucose utilization. The results showed that Fisetin treatment improved glucose clearance as measured by GTT (Figs. 2k, m) in male mice. Although D+Q treated female mice accumulated more adipose tissues they did not have impaired glucose metabolism (Figs. 2l-m). No differences in insulin sensitivity between treated and littermate control mice were observed as determined with ITT (Figs. 2n-p). Additionally, plasma insulin concentrations (Fig. 2q) were similar across treatments in both sexes of C57BL/6 mice. Plasma adiponectin (an adipokine that is involved in regulating glucose levels) concentration was significantly increased in the D+Q and Fisetin treated males, but not after either treatment in female C57BL/6 mice (Fig. 2r). The concentration of plasma insulin was elevated while adiponectin was decreased in male C57BL/6 mice compared to female littermates. This is consistent with the literature where adiponectin levels in male mice (32) and humans (33) are lower compared to females. The data indicated that monthly Fisetin treatment improved peripheral glucose metabolism in the male mice with increased adiponectin levels and enhanced glucose clearance. In female C57BL/6 mice, D+Q treatment increased WAT accumulation without affecting glucose metabolism. This may be reflective of the GTT assay being conducted in the middle rather than at the end of the experimental design.

### Fisetin treatment enhanced energy metabolism in male C57BL/6 mice, while D+Q treatment reduced energy metabolism in female C57BL/6 mice

Energy metabolism is strongly associated with aging and is a contributing factor to adipose accumulation (34, 35). In the present study, the treatment effects on energy metabolism were assessed by indirect calorimetry. Oxygen consumption (VO_2_) (Figs. 3a, c) and energy expenditure (EE) (Fig. 3d, f) were increased after Fisetin treatment in male C57BL/6 mice, while both parameters were decreased by D+Q treatment in female littermates (Figs. 3b-c, e-f). Thus, Fisetin enhanced energy metabolism in male C57BL/6 mice, while D+Q reduced energy metabolism in the female mice, potentially contributing to their WAT accumulation. Respiratory quotient (RQ) is the ratio of carbon dioxide produced and oxygen consumed by the body, reflecting substrate utilization for energy generation (36). In the current study, RQ for Fisetin treated male C57BL/6 mice was reduced indicating a shift in metabolic substrate utilization from mixed lipids and carbohydrates (RQ ∼ 0.8) to predominantly lipids (RQ ∼ 0.7) (36) (Fig. 3g, i). The RQ of female C57BL/6 mice treated with D+Q did not change (Fig. 3f). Female C57BL/6 mice had higher levels of all three measures compared male littermates. In summary, monthly Fisetin treatment in male C57BL/6 mice increased energy metabolism as indicated by enhanced VO_2_ and EE and shifted substrate utilization to predominantly lipids. In contrast, D+Q treatment decreased energy metabolism in female C57BL/6 mice as indicated by reduced VO_2_ and EE without a concomitant change in substrate utilization.

**Figure 3:**
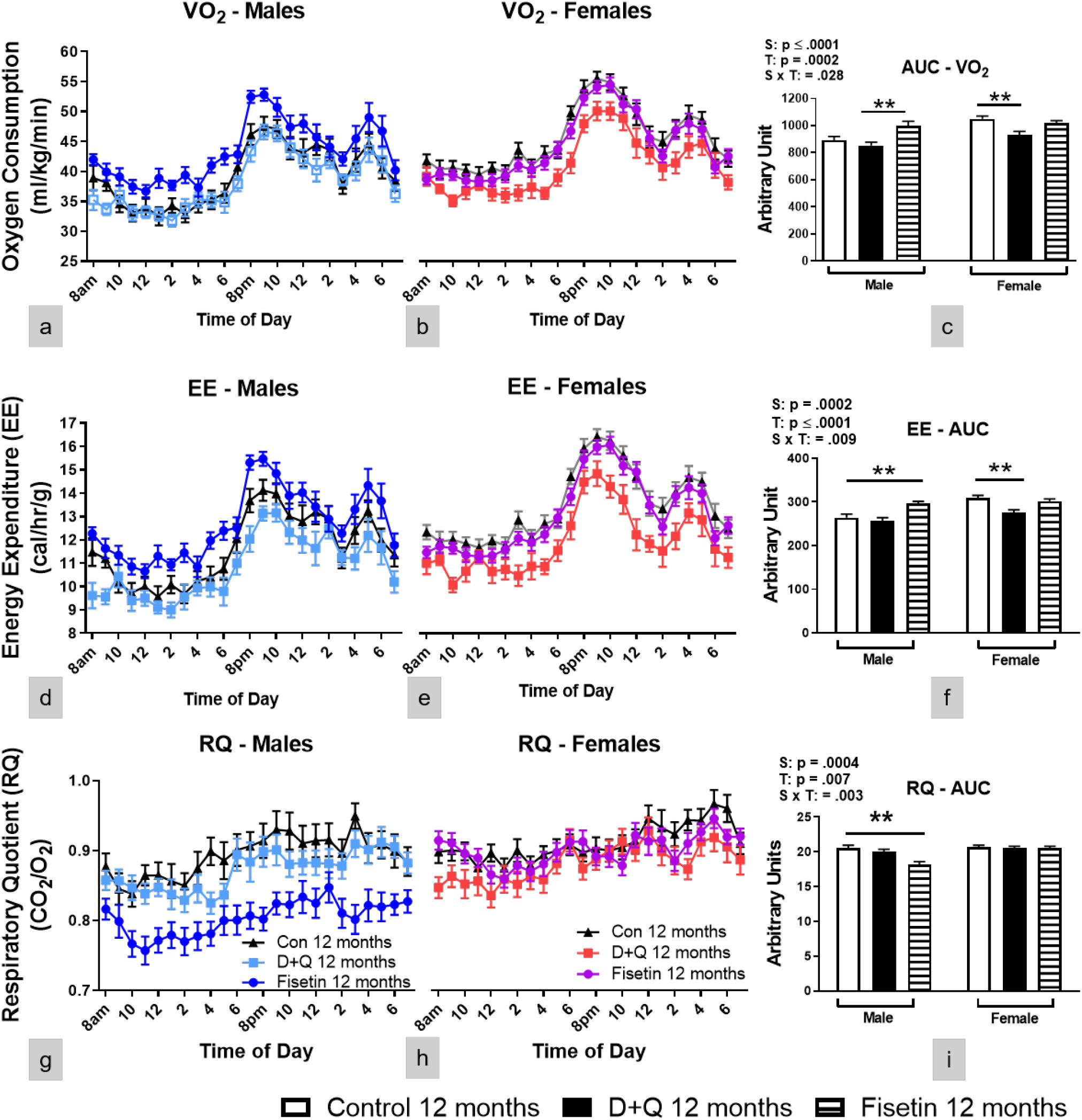
Fisetin treatment enhanced energy metabolism in male mice, while D+Q treatment decreased energy metabolism in female mice. Twenty-four-hour male and female oxygen consumption (a-c; VO_2_), energy expenditure (d-f; EE), and respiratory quotient (g-i; RQ) along with their corresponding AUC after 9 treatments. Data are presented as means ± SEM (n = 16-20). A two-way ANOVA was used to determine P-values for the categorial variables (S = Sex and T = Treatment) and their interaction (S x T), which are shown for each bar graph. **p ≤ 0.01 based on a two-tailed Student’s t test.

### Fisetin treatment improved spatial cognitive performance in male C57BL/6 mice, while D+Q treatment impaired object recognition in female C57BL/6 mice

Although deletion of senescent cells by the senolytic compounds led to functional improvements in models of neurodegenerative diseases (23, 24), it remains to be elucidated whether the monthly senolytic treatment could affect cognitive performance in normal aging mice. To investigate the effects of the treatment on cognitive functions, the MWM spatial learning and memory paradigm and a NOR task were performed. The results from MWM indicated that over 5 training sessions, the path efficiency was increased (Fig. 4a, c), and the corrected integrated path length (CIPL) (Fig. 4d, f) was reduced significantly in the Fisetin-treated male C57BL/6 mice indicative of improved learning. Neither Fisetin nor D+Q treatment affected MWM learning in the female C57BL/6 mice (Figs. 4b-c, e-f). Platform entries during the delayed probe challenge was not affected by either treatment in both sexes (Fig. 4g) suggesting spatial memory recall was unaffected. The results of NOR showed that the exploration preference for the novel object measured by retention index was decreased in the D+Q treated female mice (Fig. 4h), indicating reduced memory recall in this group. Since senotherapeutic treatment had differential effects on energy expenditure between the sexes, we wanted to verify changes to cognitive performance were not due to motor function. Mean swimming speed was similar for all treatment groups and across sexes for the training sessions and probe challenge (SI Figs. 3a-d). Additionally, no treatment differences in grip strength were observed (SI Fig. 3e). Female C57BL/6 mice had better grip strength compared to littermate males, which was consistent with other reports in C57BL/6 mice (37).

**Figure 4:**
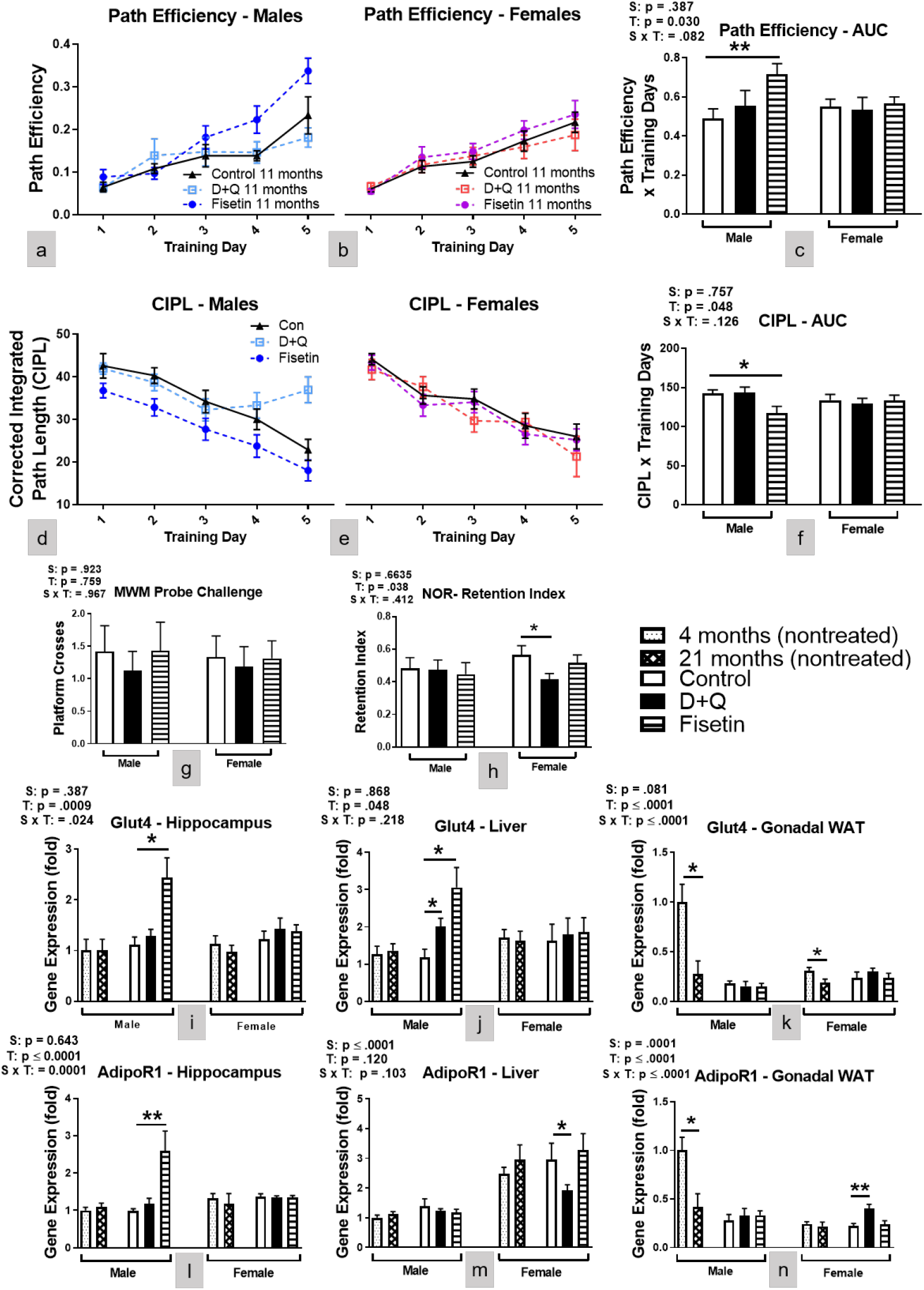
Fisetin treatment improved spatial learning in male mice, while D+Q treatment impaired object recognition memory recall in female mice. Assessment of cognition and memory recall was determined by MWM and NOR. MWM path efficiency (a-c) and corrected integrated path length (d-f; CIPL) in male and female C57BL/6 mice during 5 training sessions. Number of platform crosses during the delayed probe challenge (g). Retention index of the familiar object after a delayed novelty introduction (h). Hippocampal, liver, and gonadal WAT mRNA expression levels of Glut4 (i-k) and AdipoR1 (l-n) were measured at time of euthanization after 10 treatments. Data are presented as means ± SEM (n = 16-20). A two-way ANOVA was used to determine P-values for the categorial variables (S = Sex and T = Treatment) and their interaction (S x T), which are shown for each bar graph. *p ≤ 0.05, **p ≤ 0.01 based on a two-tailed Student’s t test.

Considering the effects of the treatments on cognitive performance, several genes related to memory formation and synaptic plasticity (38) were examined by RT-PCR in the hippocampus. Genes associated with synaptic vesicle release and glutamatergic signaling were not affected by either treatment in male or female C57BL/6 mice (SI Fig. 4). Next, we examined glucose transporter 4 (Glut4) hippocampal mRNA expression because central glucose metabolism is directly related to cognitive performance. Glut4 was increased in the hippocampus of Fisetin treated male C57BL/6 mice while D+Q had no effect in either sex (Fig. 4i). Peripherally, D+Q and Fisetin treatment increased Glut4 liver expression in male C57BL/6 mice only, but neither treatment altered expression in gonadal WAT (Figs. 4j-k). Since plasma adiponectin concentration was increased after senotherapeutic treatment in male C57BL/6 mice, we also examined adiponectin receptor 1 (AdipoR1) mRNA expression in both the hippocampus and peripheral tissues. Adiponectin signaling modulates glucose metabolism while providing anti-inflammatory effects both of which improve cognitive performance. Fisetin treated male C57BL/6 mice had increased hippocampal AdipoR1 expression, but D+Q treatment did not alter expression levels of either sex (Fig. 4l). Peripherally, neither senotherapeutic treatment affected AdipoR1 expression in the liver nor gonadal WAT of male C57BL/6 mice (Figs 4m-n). D+Q treatment reduced AdipoR1 expression in the liver while increasing expression in gonadal WAT of female C57BL/6 mice (Figs. 4m-n). This dichotomy is a result of tissue specific AdipoR1 regulation on lipid storage and utilization (39). Decreased AdipoR1 signaling in the liver increases hepatic lipogenesis causing triglyceride release and accumulation. Increased AdipoR1 expression in WAT reduces lipolysis. Together, this helps explain the increased body and WAT weight seen in C57BL/6 females mice receiving D+Q treatment. Overall, the effects of serotherapeutic treatment on Glut4 and AdipoR1 expression is consistent with changes to peripheral glucose metabolism (Figs. 2) and energy expenditure (Figs. 3).

### Comparing the rate of biological aging between sexes of C57BL/6 mice

Finally, to better understand why senotherapeutic treatment, particularly Fisetin, was only effective in males, we compared physical (SI Fig. 5a), metabolic (SI Fig. 5b-f), and cognitive (SI Fig. 5g-i) in four and thirteen-month-old C57BL/6 mice to determine potential differences in the rates of biological aging. Compared to female littermates, four-month-old mice male C57BL/6 mice have worse metabolic parameters as observed by GTT, oxygen consumption, and energy expenditure. Physical and cognitive parameters are similar between sexes at this age. By 13 months of age, an aging effect is observed whereby physical, metabolic, and cognitive parameters were reduced in both sexes of C57BL/6 mice. Since swimming speed declines with age this can influence the duration to the platform and the number of platform crosses on the MWM (SI Figs. 5g, i, j). Although the CIPL calculation avoids variability in results due to swimming speed differences, both sexes of 13-month-old C57BL/6 mice had worse spatial navigation learning and memory recall compared to their younger sex-matched littermates (SI Fig. 5h, k). These baseline differences between sexes of C57BL/6 mice may partially explain why Fisetin was effective only in males

## Discussions and conclusions

Administration of senolytic compounds is a novel approach to removing senescent cell accumulation and thereby reducing inflammation. These compounds are being investigated as potential lifespan and healthspan extending therapies and may also have applications in the treatment of numerous metabolic and neurodegenerative disorders (18, 23-25). Since Fisetin and Q are naturally occurring plant flavonoids, they are readily available as dietary supplements for use without a prescription. There is little information on the efficacy of these compounds when taken at young ages and for prolonged periods of time. The data presented here indicates that these compounds have sexually dimorphic effects, each with health benefits and risks. The monthly dosing strategy used in this study was based on previous publications showing that markers of cell senescence are still reduced after a four week off-treatment period (7). Prior research suggests that senolytic compounds work either by clearing senescent cells or reversing cellular senescence rather than preventing cells from entering a senescent state. Since senescent cell accumulation occurs over weeks, maintaining an effective concentration dosage is not required. Rather, intermittent dosing may be preferable thereby avoiding adverse effects.

Monthly Fisetin treatment was efficacious in male C57BL/6 mice when started at four months of age. This dosing strategy had little to no effect in female littermates when compared to vehicle treatment. Nine months of Fisetin treatment efficiently reduced SASP in WAT (Fig. 1), hippocampus (SI Fig. 1), and plasma (SI Fig. 2), of thirteen-month-old male C57BL/6 mice as lower mRNA expression of p16^Ink4α^, p21^Cip1^, TNFα, and IL-6 were observed. Male C57BL/6 mice receiving Fisetin treatment also had improved glucose metabolism as supported by better glucose clearance (Figs. 2k, m) after five treatments and elevated plasma adiponectin (Fig. 2r) collected upon euthanization at thirteen months of age. Improved energy metabolism was also supported by increased VO_2_ and EE (Figs. 3a, c-d, f) along with reduced RQ (Fig. 3g, i) when measured at 12 months of age. Monthly Fisetin treatment also reduced SASP markers to similar levels as four-month-old treatment naïve male C57BL/6 mice. The metabolic improvements also mirrored levels observed in younger littermates. But physical parameters (swimming speed and grip strength) as well as learning and memory recall was still worse after Fisetin treatment when compared with younger male C57BL/6 mice. This suggests administering Fisetin treatment at younger ages has a stronger anti-inflammatory (potentially mediated through increased plasma adiponectin signaling) rather than an overall anti-aging effect. This cannot be definitively concluded without frailty or lifespan measurements which were outside the scope of the study. Also, inflammation and aging are not mutually exclusive whereby chronic inflammation contributes to worse health and the onset numerous age-related ailments and neurodegenerative disorders. Approaches to reduce chronic inflammation have the potential to extend quality of life during aging.

In the present study, D+Q treatment showed minimal efficacy in male C57BL/6 mice and was detrimental to female mice. In females, SASP markers (Fig. 1) and body weight were increased attributed to larger WAT depots (Fig. 2a-e). Several factors contributed to this increased abdominal adiposity. These include increased ACC (the rate-limiting enzyme in lipid synthesis) and AdipoR1 in WAT (Fig. 2h and Fig. 4n), reduced UCP1 and PPARγ activity in BAT (Figs. 2i-j), reduced energy metabolism and liver AdipoR1 expression (Fig. 3 and Fig. 4m). Because obesity can lead to adipose tissue entering a senescent-like state at a young age (40), the accumulation of abdominal WAT in D+Q treated female mice could result in increased SASP. This possibility is supported by expression of SASP-related genes in D+Q treated females resembling or even exceeding the values measured in much older (twenty-one-month-old) untreated females (Fig. 1, SI Fig. 1, and SI Fig. 2). The reasons for the detrimental outcome of D+Q treatment in the female mice remain to be fully elucidated, but are not unique to this study. A recent publication using a similar D+Q dosing regimen reported similar observations in a C57BL/6 mouse model of hepatocellular carcinoma (41). Others have proposed that continuous senescent cell removal beginning at young ages does not always activate cell-replacement mechanisms, but rather induces regenerative responses causing fibrosis (42). A better understanding of cell-type specific effects of senescent cell removal is warranted.

Sexually dimorphic biological aging may partially explain the senotherapeutic differences observed in C57BL/6 mice. Four-month-old male and female C57BL/6 treatment naïve mice have similar SASP mRNA and plasma concentrations (Fig. 1 and SI Fig. 2), but males have worse metabolism (SI Figs. 5b, d-e). SASP became more prominent in male C57BL/6 mice compared with age-matched thirteen-month females, particularly proinflammatory cytokines TNFα and IL-6 (Fig. 1d, f and SI Fig. 2a, c). Since Fisetin shows therapeutic efficacy by reducing inflammation leading to metabolic improvement, these SASP factors were not at harmful levels in female C57BL/6 mice. This suggests male C57BL/6 mice age biologically faster and may explain why Fisetin treatment was ineffective at the ages tested in female littermates. Unfortunately, data is only beginning to emerge regarding sexually dimorphic aging characteristics due in large part from male centric sampling in both preclinical and clinical research. Sparse evidence suggests females maintain better cellular health throughout the majority of their life, but become frailer when approaching death compared with males (43).

In C57BL/6 mice, cell senescence usually becomes prominent by approximately 14 months of age (7), and treatment with senolytic drugs at this or a later ages cleared senescent cells and led to various beneficial effects (20, 21). However, it was unclear whether chronic treatment with senolytic drugs started at an earlier age (prior to reported senescent cell accumulation) in C57BL/6 mice would have similar effects as the reported observations during later age (20, 21). This gap in knowledge is particularly relevant given that Fisetin and Q are available without a prescription and may be taken by younger adults. Our data revealed that such treatment indeed had impacts on the mice as they reached median lifespan. However, the senotherapeutic treatment effects were sexually dimorphic. C57BL/6 male, not female, mice receiving Fisetin treatment had beneficial responses mediated through reduced SASP and improved glucose and energy metabolism. Female, not male, mice receiving a D+Q cocktail treatment had detrimental responses with reduced energy metabolism, increased WAT accumulation, and elevated SASP.

D+Q was the first well-characterized senotherapeutic agent proven to reduce senescent cell burden and extend lifespan. Despite Q and Fisetin being structural analogs, Q had minimal to no effects when initiated at a young age in male C57BL/6 mice. Although both are potent redox-active flavonoids, Fisetin has higher prooxidant activity. Furthermore, Q’s senolytic potency is augmented by trace elements such as copper and iron that are known to accumulate during aging and in senescent cells (5). Since senolytic drugs were administered at a young age prior to reported senescent cell accumulation, iron and copper accumulation was not a threshold to enhance Q’s potency. Fisetin’s prooxidant activities are not amplified in the presence of copper or iron, which may explain why health benefits were only observed with this senotherapeutic in male C57BL/6 mice.

Moreover, Fisetin-treated male C57BL/6 mice not only had improved markers of peripheral health, but also their spatial learning during the MWM training sessions (Figs. 4a, c, d, f) compared to vehicle treated C57BL/6 mice. These cognitive improvements, however, did not reach levels observed in 4-month-old treatment naïve mice (Fig. 4g and SI Fig. 5i). Although we did not observe changes in markers of synaptic plasticity associated with hippocampal memory formation (SI Fig. 4), we did observe elevated expression of Glut4 (Fig. 4i-j), hippocampal AdipoR1 (Fig. 4l), and plasma adiponectin concentrations (Fig. 2r) suggesting enhanced glucose transport and adiponectin signaling in the brain. This novel finding that Fisetin treatment concurrently improved peripheral as well as central glucose and adiponectin signaling has important implications for metabolic and cognitive function. In the periphery, adiponectin increases insulin sensitivity, stimulates fatty acid oxidation and glucose uptake, suppresses hepatic glucose production (44), and inhibits inflammation (45). In the brain, adiponectin signaling via AdipoR1 has anti-inflammatory effects and is involved with cognitive and neuroprotective mechanisms (46). Monthly Fisetin treatment in male mice led to increased adiponectin levels in circulation, which likely contributed to improvements of peripheral glucose and lipid metabolism. Circulating adiponectin passes through the blood brain barrier where it can enhance cognitive function by upregulating glucose transport and AdipoR1 signaling (46). However, neither hippocampal Glut4 (Fig. 4i), AdipoR1 (Fig. 4l), nor the examined synaptic plasticity markers (SI Fig. 4) were affected by D+Q treatment in the female mice even though object recognition memory retention was reduced (Fig. 4h). The reduced cognitive performance of these animals might be attributed to the increased adiposity after D+Q treatment, but future studies are needed to identify the mechanisms involved.

Cell senescence is strongly correlated with aging, but currently no diagnostic testing is available to quantify senescent cells accumulation. It seems possible that the earlier in life senotherapeutics are taken, the better likelihood of delaying age-related decline and disease risk might be expected. Senolytics such as Fisetin and D+Q are known to reduce senescent cell burden (7), but the optimal age to begin treatment is currently unknown. Our study indicates that both biological aging and sex may also determine the therapeutic outcomes of senolytic treatment. When the treatment was started at 4 months of age, before the reported senescent cell accumulation, Fisetin had beneficial effects in male C57BL/6 mice while a D+Q cocktail had adverse consequences in female C57BL/6 mice.

These observations provide novel information with translational relevance. First, senolytic drugs taken at an age before significant senescent cell burden to reduce or prevent their prevalence later in life may be detrimental to overall health. Second, males and females can have differential responses to the same senolytic treatment when initiated at younger ages. Third, a particular senolytic treatment may have beneficial, negligible or detrimental effects depending on the age, sex, or disease state. Fourth, oral senolytic treatment not only impacted peripheral physiological responses, but also affected cognitive performance, possibly via increased central adiponectin signaling and glucose transport. Overall, these observations should serve as a note of caution in this rapidly evolving and expanding field of investigation.

Designing senotherapeutics with the role of sex and age of onset could conceivably provide a useful preventative treatment strategy for dealing with development of neurodegenerative disorders. Supporting this possibility, development of metabolic syndrome in midlife was reported to increases the risk for Alzheimer’s disease (AD) later on (47). Besides clearance of senescent cells, senotherapeutics may ameliorate or delay cognitive decline through mechanisms associated with improved peripheral and central glucose metabolism. Considering metabolic dysregulation has been observed in the APP/PS1 model of AD (48), Additional studies are warranted to determine if prodromal senolytic treatment could provide cognitive benefits in neurodegenerative diseases.

## Material and methods

### Chemicals

Unless otherwise noted, all chemicals were obtained from Sigma-Aldrich (St. Louis, MO) including Quercetin (Cat# RHR1488). Fisetin was purchased from Selleckchem (Houston, TX; Cat #S2298), and Dasatinib from LC laboratories (Woburn, MA; Cat# D-3307).

### Animals

Male and female C57BL/6 mice were maintained in our established breeding colonies. Protocols for animal use were approved by the Institutional Animal Care and Use Committee at Southern Illinois University School of Medicine. Mice were group housed on a 12:12 h light-dark cycle, and food (Chow 5001 with 23.4% protein, 5% fat, and 5.8% crude fiber; LabDiet PMI Feeds) and water were available ad libitum. Sentinel mice located in the same rooms as our breeder and experimental mice were tested triannually for pyrogens and pathogens and none were detected. All assays, including tissue collection, were conducted one week after monthly senotherapeutic dosing. As shown in Fig 1a, glucose metabolism was assessed via ITT and GTT after six treatments at the animal age of nine months. Grip Strength was measured after seven treatments while NOR and MWM was performed after eight treatments. Energy metabolism was measured by indirect calorimetry at the animal age of twelve. A week following the final senotherapeutic treatment at 13 months, body weight was recorded then euthanized with an overdose of isoflurane. A cardiac puncture was used to collect blood for assessment of circulating SASP markers and proteins. Mice were rapidly decapitated and the peripheral tissues were weighed to calculate their contribution to total body weight. Tissue was immediately flash frozen and stored at -80°C until processing as previously described (49).

### Senolytic drug treatment

Senotherapeutic concentrations and dosing strategy were based on previous publications (20, 50-52). C57BL/6 mice were dosed with 100 mg/kg of Fisetin while a cocktail of 5 mg/kg of D + 50 mg/kg of Q, or vehicle (2% DMSO in canola oil) by oral administration (53). The treatments were given once per month from 4-13 months of age.

### Glucose tolerance test (GTT) and insulin tolerance test (ITT)

GTT or ITT was carried out as described previously (49). Sixteen-hour-fasted mice underwent GTT by intraperitoneal (i.p.) injection with 1 g glucose per kg of body weight (BW). Blood glucose levels were measured at 0, 15, 30, 45, 60, and 120 min with a PRESTO glucometer (AgaMatrix). For ITT, nonfasted mice were injected i.p. with 1 IU porcine insulin from sigma (St. Louis, MO) (Cat# I5523) per kg of BW. Blood glucose levels were measured at 0, 15, 30, 45, 60, and 120 min. The data for GTT are presented as absolute value, and for ITT are presented as a percentage of baseline glucose.

### Indirect calorimetry

Indirect calorimetry was performed as previously described (49) using AccuScan Metabolic System (AccuScan Instruments). In this system, mice are housed individually in metabolic chambers with ad libitum access to food and water. After a twenty-four-hour acclimation period, VO_2_, VCO_2_, EE and RQ measurements were collected every ten min per animal and averaged for each hour.

### Morris water maze (MWM) training and probe challenge

The MWM was used to assess spatial learning and memory recall, and performed as previously described (54). Mice were trained to utilize visual cues placed around the room to repeatedly swim to a static, hidden escape platform (submerged one cm below the opaque water surface), regardless of starting quadrant. The MWM paradigm consisted of 5 consecutive training days with three, 90 s trials/day and a minimum inter-trial-interval of 20 min. Starting quadrant was randomized for each trial. After two days without testing, the escape platform was removed and all mice entered the pool of water from the same starting position for a single 60 s probe challenge to test long-term memory recall. The ANY-maze video tracking system (Stoelting Co., Wood Dale, IL) was used to record mouse navigation during the training and probe challenge. The three trials for each training day were averaged for each mouse.

### Novel Object Recognition (NOR)

The NOR was used to evaluate memory retention based on the premise that mice spend more time exploring a novel object rather than a familiar object if memory capabilities remain intact. Mice were habituated in the open field chamber for 30 min on the first day. Twenty-four hours after, the mouse was returned to the chamber and presented with two similar objects for 5 min. A 24-hour inter-session-interval was used between introduction and retention phases to assess long-term memory retrieval. During the retention phase one of the familiar objects is replaced with a novel object and the mouse is given 5 minutes of exploration. The ANY-maze video tracking system (Stoelting Co., Wood Dale, IL) was used to record mouse navigation during the familiarization and retention phases and time spent at each object. The time spent exploring the novel object was divided by the total time spent exploring both objects to calculate the retention index.

### Grip Strength

Using their forepaws, mice grasp a wire grid connected to an isometric force transducer (Harvard Apparatus; Holliston, MA) and are gently pulled horizontally away by the tail. Force is measured in grams and normalized according to body weight. Three trials with a minimum inter-trial-interval of twenty minutes are averaged per mouse.

### Assessment of blood chemistry

Plasma was collected from animals anesthetized with isoflurane by cardiac puncture at sacrifice. The blood was mixed with EDTA, followed by centrifugation at 10,000 g for 15 min at 4°C for plasma collection. Per the manufacturer’s protocol, insulin or adiponectin was measured with respective ELISA kits (Crystal Chem, Elk Grove Village, IL; Cat# 90080 and 80569). Plasma cytokine levels were assayed using a multiplex immunoassay on a MESO QuickPlex SQ 120 with accompanying software (Meso Scale Diagnostics, LLC).

### RT–PCR

mRNA expression was analyzed by quantitative RT–PCR as previously described (49) using cDNA synthesis kits and SYBR green mix from Bio-Rad (Cat# 1708897 and 1725121) performed with the StepOne Real-Time PCR System (Thermo Fisher Scientific). RNA was extracted using an RNeasy mini kit or RNeasy Lipid Tissue Mini Kit (Qiagen) following the manufacturer’s instructions. Relative expression was calculated as previously described and primers were purchased from Integrated DNA Technologies (Supplemental Information Table 1).

### Statistical analysis

Statistical analyses were conducted using a two-way ANOVA to test for significance of sex (males × females), treatment (Vehicle × Fisetin × D+Q), and an interaction between sex and treatment. Difference between two groups were assessed with unpaired two-tailed Student’s t tests with significance defined as p ≤ 0.05. The area under the curve was calculated using the two-way ANOVA. Data are presented as means ± SEM. All statistical analyses and graphs were completed using Prism 9 (GraphPad Inc, La Jolla, CA, USA).

## Acknowledgements

This work was supported by the National Institutes of Health NIA R01-AG057767 and NIA R01-AG061937, Dale and Deborah Smith Center for Alzheimer’s Research and Treatment, Kenneth Stark Endowment (YF, SF, KQ, MRP, KNH, ERH), NIA R21-AG062985, and American Diabetes Association 1-19-IBS-126 (DM, RS, AB). We would like to thank Melissa Roberts for performing and analyzing the plasma cytokine multiplex assay and Lisa Hensley for editorial support.

## Author Contributions

YF, DM, RS, SM, KQ, and MRP conducted the experiments and analyzed the data. YF, AB, KNH, and ERH conceived the study, designed the experiments, interpreted the data, and wrote the manuscript. All authors approved the final version of the manuscript.

## Data Availability

All data is available upon reasonable request.

## Data Citation

All data was generated by the authors. No external sources of data were used to create any figures presented in this manuscript.

## Conflict of Interest

The authors declare no conflicts of interest.

---

**Supplemental Information: Figure 1:**
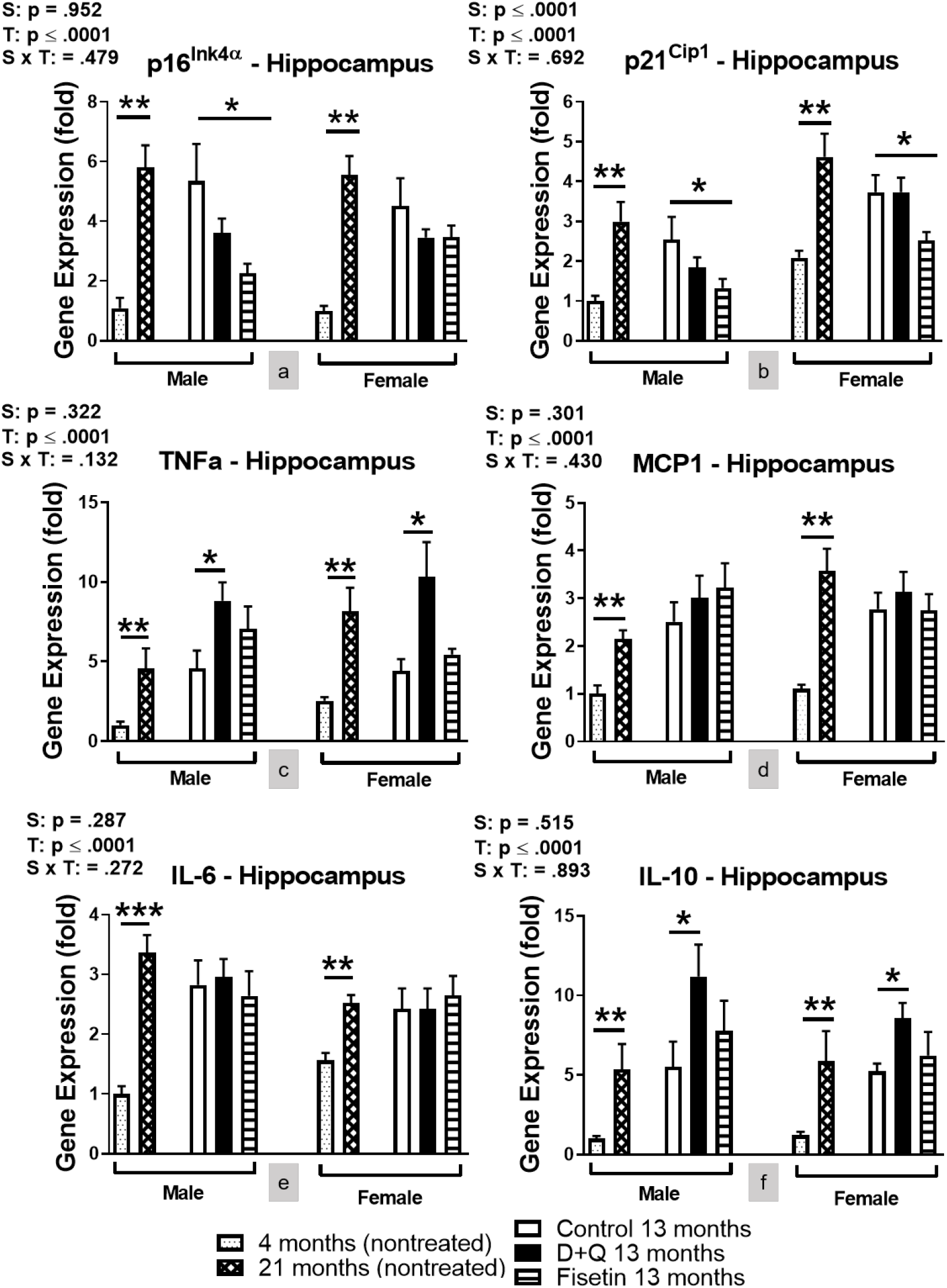
Effects of senotherapeutic treatment on the hippocampal SASP profile in C57BL/6 mice. mRNA expression levels of hippocampal SASP markers (a-f) were measured at time of euthanization after 10 treatments. Data are presented as means ± SEM (n = 8-12). A two-way ANOVA was used to determine P-values for the categorial variables (S = Sex and T = Treatment) and their interaction (S x T), which are shown for each bar graph. *p ≤ 0.05, **p ≤ 0.01, ***p<0.001 based on a two-tailed Student’s t test.

**Supplemental Information Figure 2:**
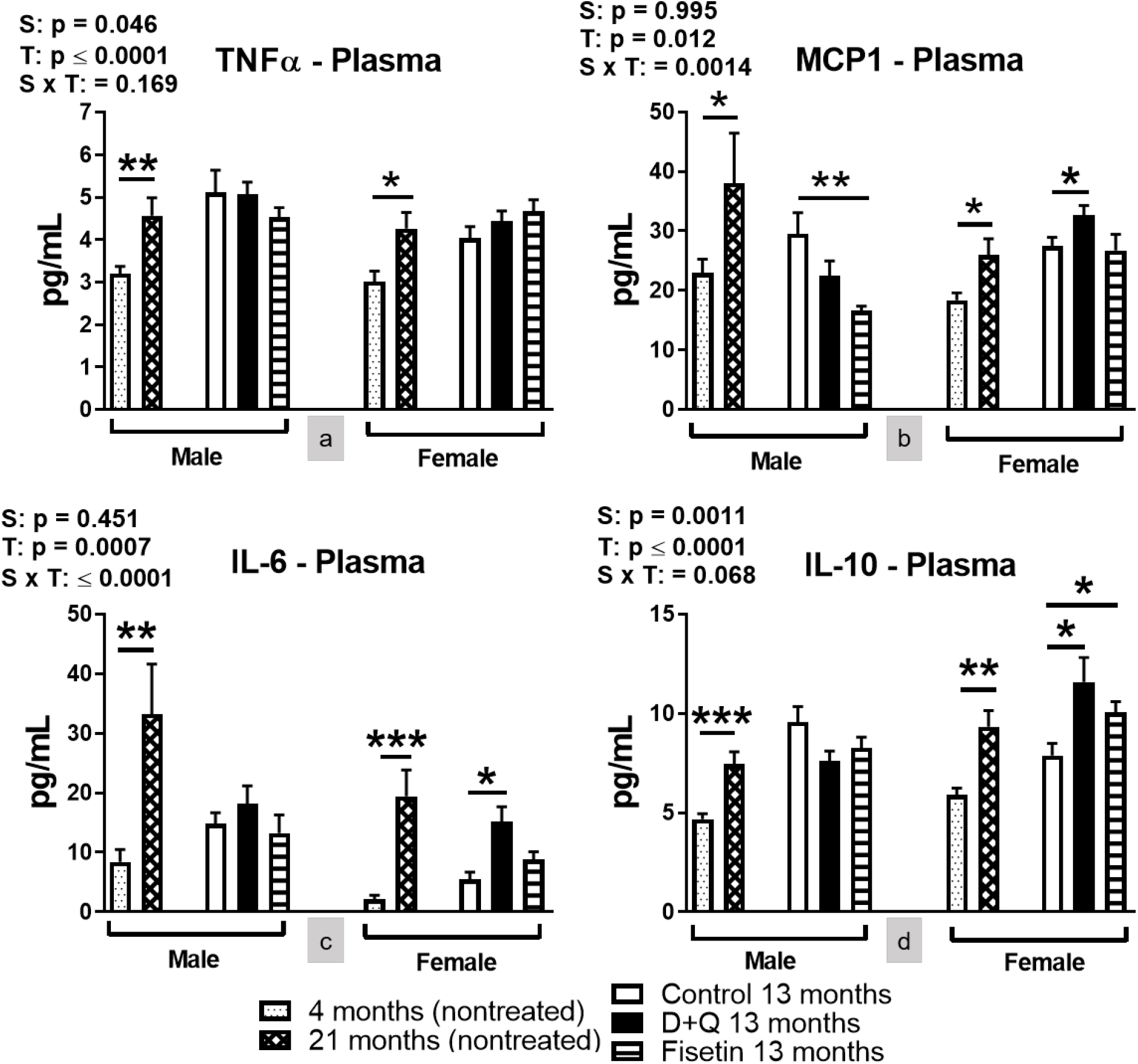
Senotherapeutic treatment altered plasma cytokine concentrations in C57BL/6 mice. Plasma concentrations of TNFα, MCP1, IL-6, and IL-10 (a-d) from time of euthanization after 10 senotherapeutic treatments. Data are represented as means ± SEM (n = 8). A two-way ANOVA was used to determine P-values for the categorial variables (S = Sex and T = Treatment) and their interaction (S x T), which are shown for each bar graph. *p ≤ 0.05, **p ≤ 0.01, ***p<0.001 based on a two-tailed Student’s t test.

**Supplemental Information Figure 3:**
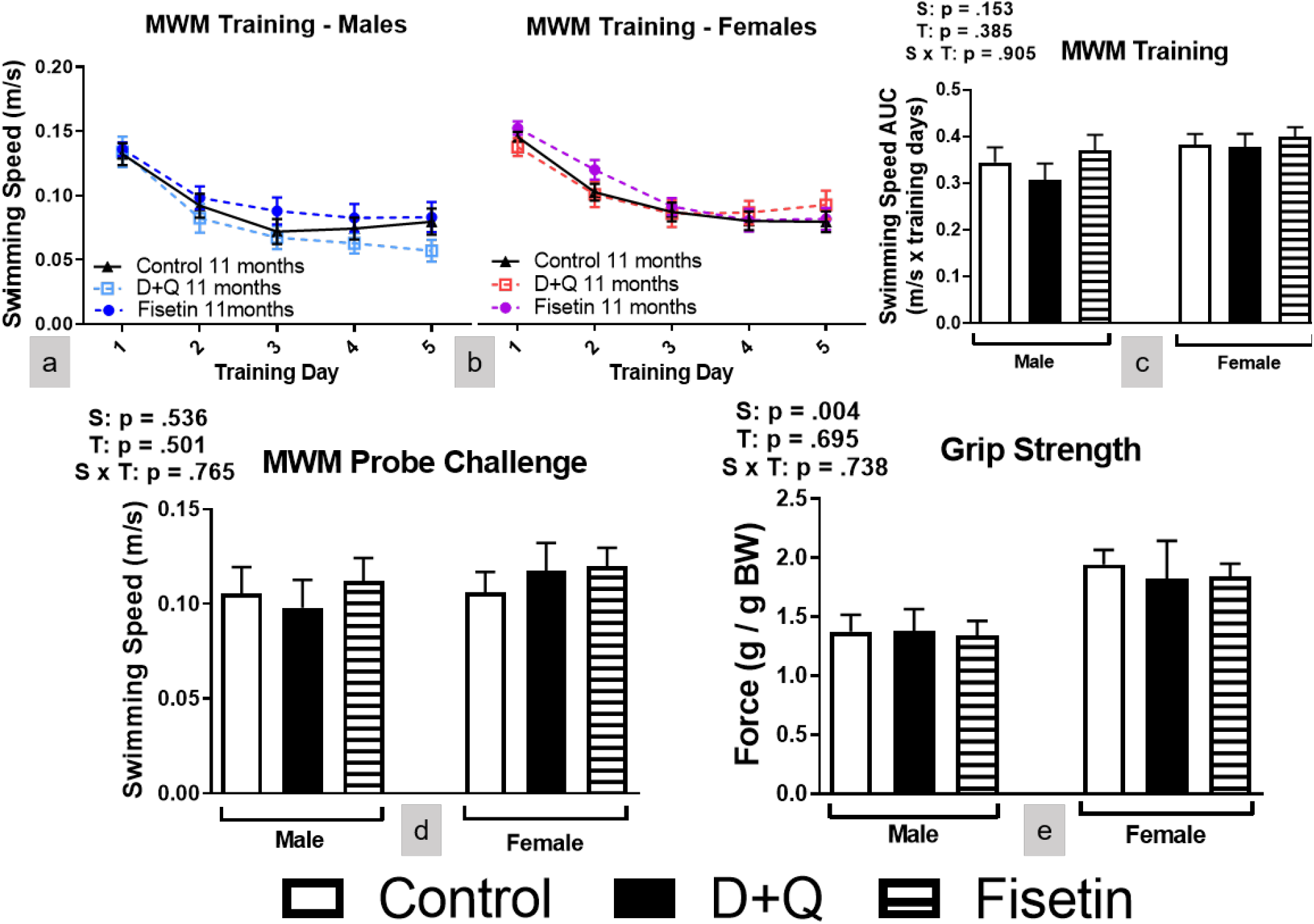
Effects of senotherapeutic treatment on physical performance in C57BL/6 mice. MWM swimming speed during the training days and probe challenge (a-d) and forepaw grip strength (e). Data are presented as means ± SEM (n = 16-20). A two-way ANOVA was used to determine P-values for the categorial variables (S = Sex and T = Treatment) and their interaction (S x T), which are shown for each bar graph.

**Supplemental Information Figure 4:**
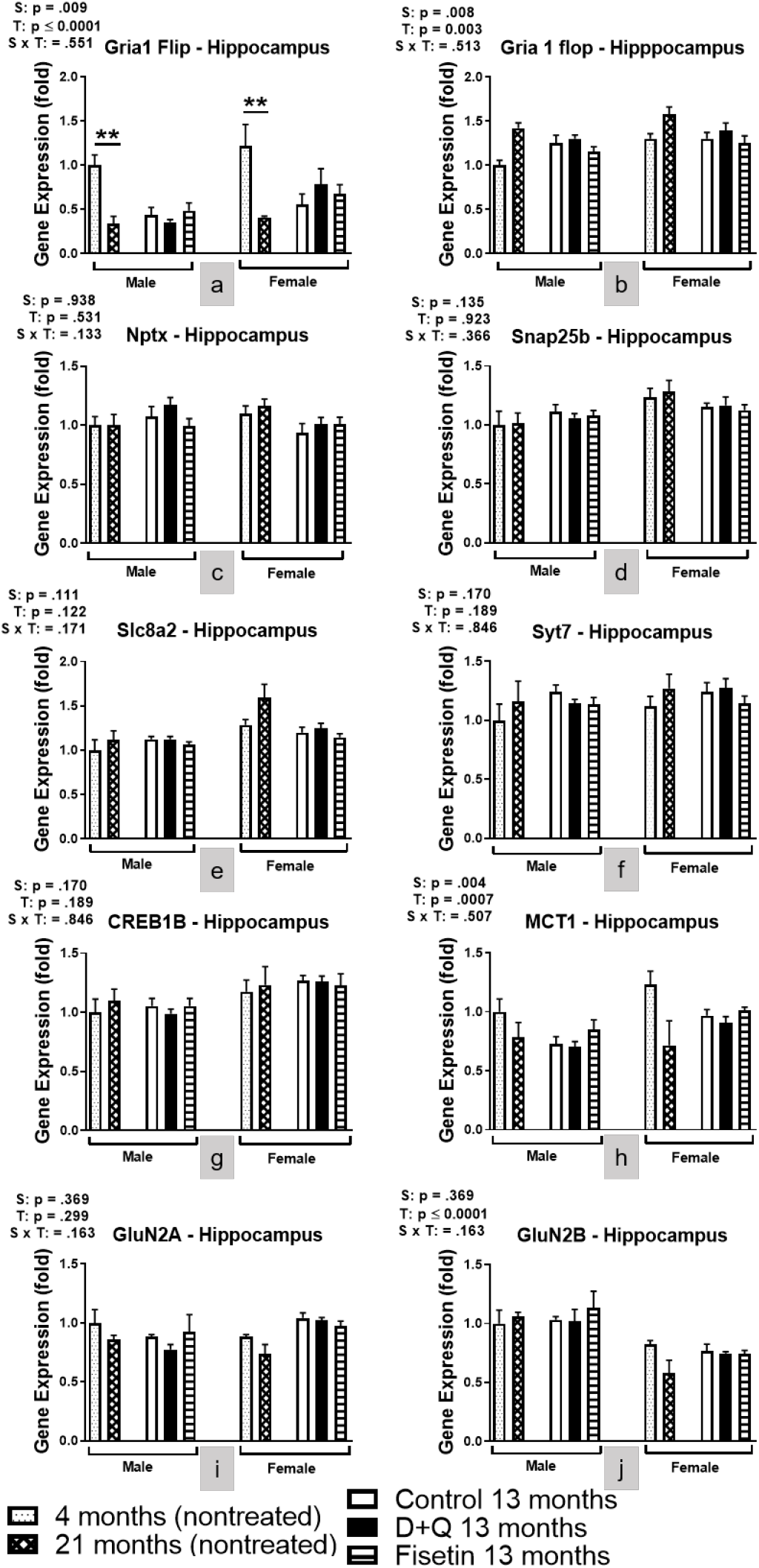
Hippocampal mRNA expression of genes involved with synaptic plasticity were unaltered after senotherapeutic treatment. Hippocampal mRNA expression from time of euthanization after 10 senotherapeutic treatments in C57BL/6 mice. Data are represented as means ± SEM (n = 16-20). A two-way ANOVA was used to determine P-values for the categorial variables (S = Sex and T = Treatment) and their interaction (S x T), which are shown for each bar graph. **p ≤ 0.01 based on a two-tailed Student’s t test.

**Supplemental Information Figure 5:**
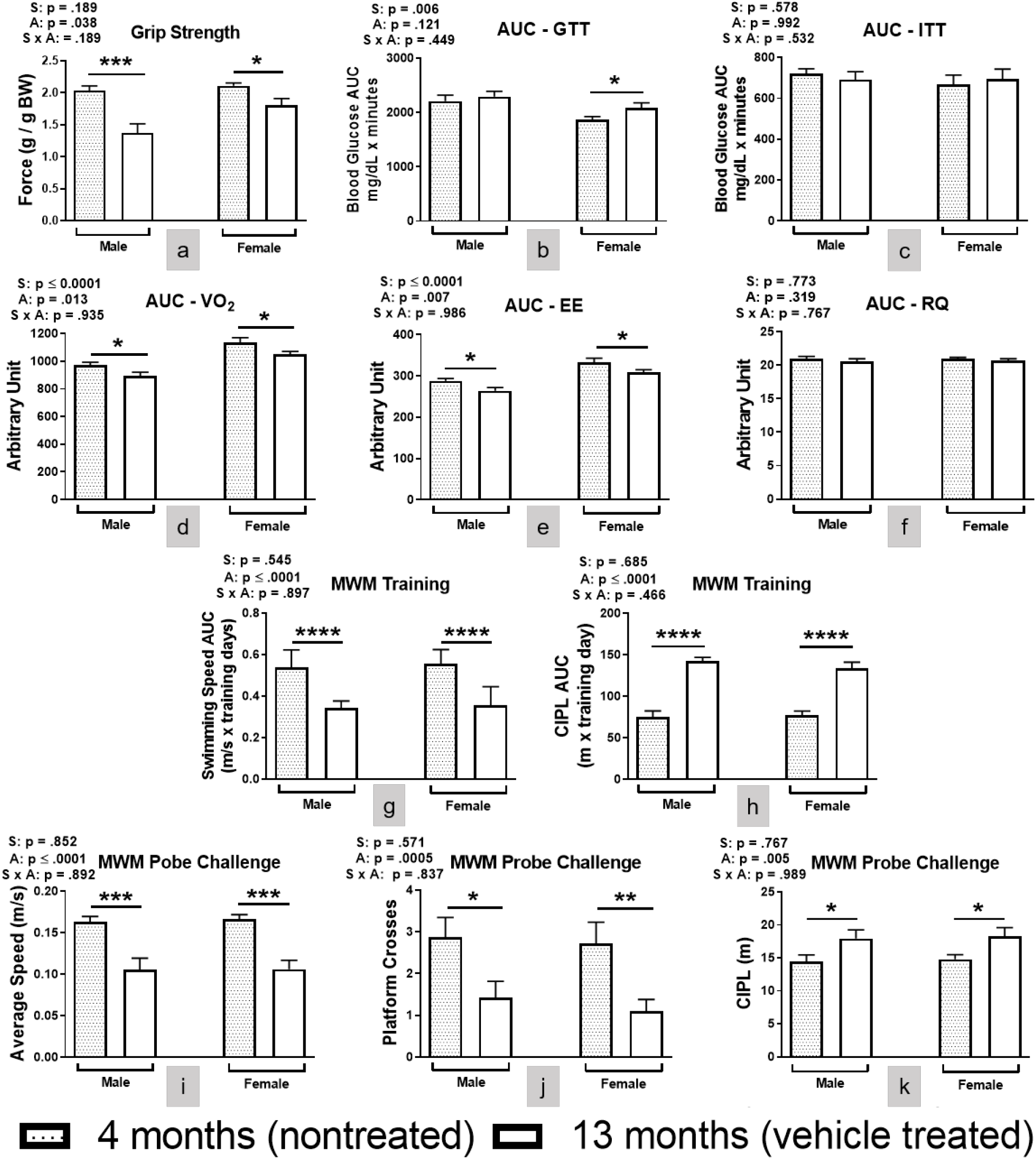
Effects of aging on physical, metabolic, and cognitive parameters. Nontreated four-month-old C57BL/6 mice were compared to sex-matched control treated mice. Data are represented as means ± SEM (n = 16-20). A two-way ANOVA was used to determine P-values for the categorial variables (S = Sex and A = Age) and their interaction (S x A), which are shown for each bar graph. *p≤ 0.05, **p ≤ 0.01, ***p<0.001 based on a two-tailed Student’s t test.

**Supplemental Information: Table 1:**
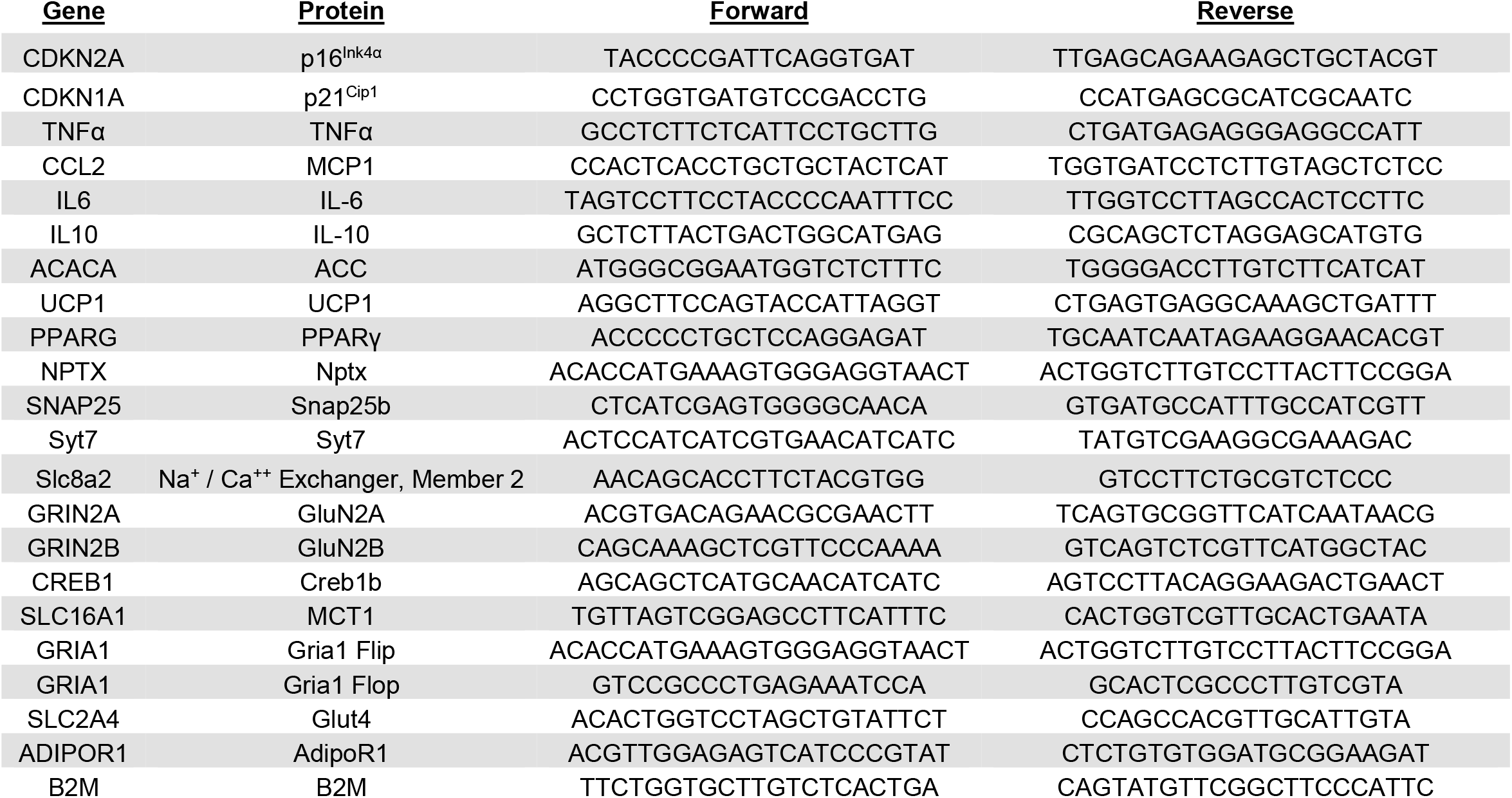
A list of forward and reverse primers used in this study.

## Notes

### Competing Interest Statement

The authors have declared no competing interest.

### Summary of Updates

Based on reviewer feedback, we have included additional experiments to the original study design. This has led to significant revisions to all sections of the manuscript.

## Reference

1. Grynkiewicz G & Demchuk OM (2019) New Perspectives for Fisetin. Front Chem 7:697.

2. Jafarinia M, et al. (2020) Quercetin with the potential effect on allergic diseases. Allergy Asthma Clin Immunol 16:36.

3. Mlcek J, Jurikova T, Skrovankova S, & Sochor J (2016) Quercetin and Its Anti-Allergic Immune Response. Molecules 21(5).

4. Aguilera DG & Tsimberidou AM (2009) Dasatinib in chronic myeloid leukemia: a review. Ther Clin Risk Manag 5(2):281–289.

5. Wang Y, He Y, Rayman MP, & Zhang J (2021) Prospective Selective Mechanism of Emerging Senolytic Agents Derived from Flavonoids. J Agric Food Chem 69(42):12418–12423.

6. Zhu Y, et al. (2015) The Achilles’ heel of senescent cells: from transcriptome to senolytic drugs. Aging Cell 14(4):644–658.

7. Kirkland JL & Tchkonia T (2020) Senolytic drugs: from discovery to translation. J Intern Med 288(5):518–536.

8. Campisi J (2005) Senescent cells, tumor suppression, and organismal aging: good citizens, bad neighbors. Cell 120(4):513–522.

9. Kuilman T & Peeper DS (2009) Senescence-messaging secretome: SMS-ing cellular stress. Nat Rev Cancer 9(2):81–94.

10. Coppe JP, Desprez PY, Krtolica A, & Campisi J (2010) The senescence-associated secretory phenotype: the dark side of tumor suppression. Annu Rev Pathol 5:99–118.

11. Lasry A & Ben-Neriah Y (2015) Senescence-associated inflammatory responses: aging and cancer perspectives. Trends Immunol 36(4):217–228.

12. Freund A, Orjalo AV, Desprez PY, & Campisi J (2010) Inflammatory networks during cellular senescence: causes and consequences. Trends Mol Med 16(5):238–246.

13. McHugh D & Gil J (2018) Senescence and aging: Causes, consequences, and therapeutic avenues. J Cell Biol 217(1):65–77.

14. Chen JH, Hales CN, & Ozanne SE (2007) DNA damage, cellular senescence and organismal ageing: causal or correlative? Nucleic Acids Res 35(22):7417–7428.

15. Chapman J, Fielder E, & Passos JF (2019) Mitochondrial dysfunction and cell senescence: deciphering a complex relationship. FEBS Lett 593(13):1566–1579.

16. Prata L, Ovsyannikova IG, Tchkonia T, & Kirkland JL (2018) Senescent cell clearance by the immune system: Emerging therapeutic opportunities. Semin Immunol 40:101275.

17. Passos JF, Miwa S, & von Zglinicki T (2013) Measuring reactive oxygen species in senescent cells. Methods Mol Biol 965:253–263.

18. Musi N, et al. (2018) Tau protein aggregation is associated with cellular senescence in the brain. Aging Cell 17(6):e12840.

19. Ogrodnik M, et al. (2019) Obesity-Induced Cellular Senescence Drives Anxiety and Impairs Neurogenesis. Cell Metab 29(5):1233.

20. Yousefzadeh MJ, et al. (2018) Fisetin is a senotherapeutic that extends health and lifespan. EBioMedicine 36:18–28.

21. Xu M, et al. (2018) Senolytics improve physical function and increase lifespan in old age. Nat Med 24(8):1246–1256.

22. Baker DJ, et al. (2011) Clearance of p16Ink4a-positive senescent cells delays ageing-associated disorders. Nature 479(7372):232–236.

23. Bussian TJ, et al. (2018) Clearance of senescent glial cells prevents tau-dependent pathology and cognitive decline. Nature 562(7728):578–582.

24. Zhang P, et al. (2019) Senolytic therapy alleviates Abeta-associated oligodendrocyte progenitor cell senescence and cognitive deficits in an Alzheimer’s disease model. Nat Neurosci 22(5):719–728.

25. Khatoon S, Agarwal NB, Samim M, & Alam O (2021) Neuroprotective Effect of Fisetin Through Suppression of IL-1R/TLR Axis and Apoptosis in Pentylenetetrazole-Induced Kindling in Mice. Front Neurol 12:689069.

26. Huang YH, et al. (2020) Interleukin-10 induces senescence of activated hepatic stellate cells via STAT3-p53 pathway to attenuate liver fibrosis. Cell Signal 66:109445.

27. Maciel-Baron LA, et al. (2016) Senescence associated secretory phenotype profile from primary lung mice fibroblasts depends on the senescence induction stimuli. Age (Dordr) 38(1):26.

28. Ou MY, Zhang H, Tan PC, Zhou SB, & Li QF (2022) Adipose tissue aging: mechanisms and therapeutic implications. Cell Death Dis 13(4):300.

29. Hudgins AD, et al. (2018) Age- and Tissue-Specific Expression of Senescence Biomarkers in Mice. Front Genet 9:59.

30. Cannon B & Nedergaard J (2004) Brown adipose tissue: function and physiological significance. Physiol Rev 84(1):277–359.

31. Kalinovich AV, de Jong JM, Cannon B, & Nedergaard J (2017) UCP1 in adipose tissues: two steps to full browning. Biochimie 134:127–137.

32. Gui Y, Silha JV, & Murphy LJ (2004) Sexual dimorphism and regulation of resistin, adiponectin, and leptin expression in the mouse. Obes Res 12(9):1481–1491.

33. Isobe T, et al. (2005) Influence of gender, age and renal function on plasma adiponectin level: the Tanno and Sobetsu study. Eur J Endocrinol 153(1):91–98.

34. Azzu V & Valencak TG (2017) Energy Metabolism and Ageing in the Mouse: A Mini-Review. Gerontology 63(4):327–336.

35. Westbrook R, Bonkowski MS, Strader AD, & Bartke A (2009) Alterations in oxygen consumption, respiratory quotient, and heat production in long-lived GHRKO and Ames dwarf mice, and short-lived bGH transgenic mice. J Gerontol A Biol Sci Med Sci 64(4):443–451.

36. Prentice RL, et al. (2013) An exploratory study of respiratory quotient calibration and association with postmenopausal breast cancer. Cancer Epidemiol Biomarkers Prev 22(12):2374–2383.

37. Baumann CW, Kwak D, & Thompson LV (2019) Sex-specific components of frailty in C57BL/6 mice. Aging (Albany NY) 11(14):5206–5214.

38. Reshetnikov VV, et al. (2020) Genes associated with cognitive performance in the Morris water maze: an RNA-seq study. Sci Rep 10(1):22078.

39. Lee B & Shao J (2014) Adiponectin and energy homeostasis. Rev Endocr Metab Disord 15(2):149–156.

40. Tchkonia T, et al. (2010) Fat tissue, aging, and cellular senescence. Aging Cell 9(5):667–684.

41. Raffaele M, et al. (2021) Mild exacerbation of obesity- and age-dependent liver disease progression by senolytic cocktail dasatinib + quercetin. Cell Commun Signal 19(1):44.

42. Grosse L, et al. (2020) Defined p16(High) Senescent Cell Types Are Indispensable for Mouse Healthspan. Cell Metab 32(1):87–99 e86.

43. Hagg S & Jylhava J (2021) Sex differences in biological aging with a focus on human studies. Elife 10.

44. Roy B & Palaniyandi SS (2021) Tissue-specific role and associated downstream signaling pathways of adiponectin. Cell Biosci 11(1):77.

45. Bloemer J, et al. (2018) Role of Adiponectin in Central Nervous System Disorders. Neural Plast 2018:4593530.

46. Rastegar S, et al. (2019) Expression of adiponectin receptors in the brain of adult zebrafish and mouse: Links with neurogenic niches and brain repair. J Comp Neurol 527(14):2317–2333.

47. Baumgart M, et al. (2015) Summary of the evidence on modifiable risk factors for cognitive decline and dementia: A population-based perspective. Alzheimers Dement 11(6):718–726.

48. Hascup ER, et al. (2019) Diet-induced insulin resistance elevates hippocampal glutamate as well as VGLUT1 and GFAP expression in AbetaPP/PS1 mice. J Neurochem 148(2):219–237.

49. Fang Y, et al. (2020) Lifespan of long-lived growth hormone receptor knockout mice was not normalized by housing at 30 degrees C since weaning. Aging Cell 19(5):e13123.

50. Palmer AK, et al. (2015) Cellular Senescence in Type 2 Diabetes: A Therapeutic Opportunity. Diabetes 64(7):2289–2298.

51. Saccon TD, et al. (2021) Senolytic Combination of Dasatinib and Quercetin Alleviates Intestinal Senescence and Inflammation and Modulates the Gut Microbiome in Aged Mice. J Gerontol A Biol Sci Med Sci 76(11):1895–1905.

52. Roos CM, et al. (2016) Chronic senolytic treatment alleviates established vasomotor dysfunction in aged or atherosclerotic mice. Aging Cell 15(5):973–977.

53. Dalterio S & Bartke A (1979) Perinatal exposure to cannabinoids alters male reproductive function in mice. Science 205(4413):1420–1422.

54. Hascup KN, Findley CA, Sime LN, & Hascup ER (2020) Hippocampal alterations in glutamatergic signaling during amyloid progression in AbetaPP/PS1 mice. Sci Rep 10(1):14503.

